# Effects of the Neutral CB1 Receptor Antagonist AM6527 on Spontaneous, Consummatory, and Motivated Behavior in Mice

**DOI:** 10.1101/2025.09.22.677777

**Authors:** Alev Ecevitoglu, Quilla C. Flanagan-Burt, Matthew Zhang, Byung Kwon Moon, Lipin Ji, Alexandros Makriyannis, Junghyup Suh

## Abstract

**Rationale:** The cannabinoid type-1 receptor (CB1R) signaling pathway plays a central role in regulating motivational and feeding behaviors. Neutral CB1R antagonists represent a promising therapeutic class with potentially fewer adverse effects than inverse agonists, yet their behavioral effects remain incompletely characterized.

**Objectives:** We investigated the behavioral profile of AM6527, orally bioavailable neutral CB1R antagonist, across naturalistic and operant paradigms in male mice. To evaluate dopaminergic involvement in AM6527’s effects, we employed several pharmacological interventions.

**Results:** Using machine learning-based Motion-Sequencing (MoSeq), which parses spontaneous behavior into sub-second syllables, we found that AM6527 did not affect overall speed in an open field, however, it increased the self-directed behaviors and reduced specific locomotor syllables at the highest dose tested. In a naturalistic reward consumption paradigm, AM6527 produced a dose-dependent reduction in milk intake. Operant conditioning paradigms revealed robust suppression of motivated responding on fixed ratio-3 and progressive ratio (PR) schedules for palatable milk reward, with the greatest impact on high-baseline performers under PR conditions. To understand the dopaminergic involvement, we co-administered dopaminergic drugs (targeting D1R, D2R, or dopamine transporter) which resulted in partial rescue of operant responding, indicating dopaminergic and non-dopaminergic contributions to AM6527’s observed behavioral effects.

**Conclusion:** Our findings suggest that neutral CB1R antagonism suppresses consummatory and motivated behaviors via dopamine-dependent and -independent mechanisms. By leveraging sub-second behavioral analysis with MoSeq, we further reveal distinct changes in spontaneous behavior, underscoring the relevance of CB-based treatments for maladaptive appetitive and motivational states in both psychiatric and metabolic disorder.

## Introduction

Cannabinoid CB1 receptors (CB1R) are broadly distributed in the brain, including the ventral tegmental area, hypothalamus, nucleus accumbens, and amygdala, and are crucial regulators of feeding, reward-seeking, and motivational behaviors (Chambers et al., 2007; Covey & Yocky, 2021; McLaughlin et al., 2005, 2006; Salamone et al., 2007; Sink et al., 2010). Early clinical studies showed that CB1R partial agonist delta 9-Tetrahydrocannabinol (THC) increases hunger, food consumption, and caloric intake (Foltin et al., 1986, 1988). Conversely, CB1R antagonists/inverse agonists such as rimonabant (SR141716A) have been shown to suppress food intake (Boyd & Fremming, 2005), leading to considerable interest in their potential as treatments for obesity and weight loss.

Preclinical studies have mirrored these findings, demonstrating that CB1R antagonists and inverse agonists suppress food intake and operant responding for palatable food. For example, rimonabant (SR141716A) has been shown to induce weight loss, suppress food consumption and appetite, and reduce lever pressing performance for food in rodents (Arnone et al., 1997; Colombo et al., 1998; Freedland et al., 2000). Despite their initial clinical relevance, CB1R inverse agonists including rimonabant have been taken off the market due to adverse side effects, including nausea and psychiatric symptoms such as depression (Boyd & Fremming, 2005; Christensen et al., 2007). This has shifted research attention toward neutral CB1R antagonists, which may offer safer profiles compared to inverse agonists. Similar to studies with inverse agonists, CB1R antagonists AM4113, AM6545, and AM6527 have been shown to dose-dependently reduce food-reinforced lever pressing under fixed-ratio (FR) schedules, such as FR-5. These compounds also decreased intake of high-fat, high-carbohydrate, and standard laboratory chow in male rats, underscoring their appetite suppressing properties (Randall et al., 2010; Sink et al., 2008, 2009). Among these, AM6527 stands out due to its greater potency and improved oral bioavailability (Sink et al., 2009). Moreover, these suppressive effects are comparable to those observed with prefeeding and aversive manipulations such as quinine adulteration (Thompson et al., 2016), suggesting that CB1R antagonist AM6527 may modulate both motivational as well as consummatory aspects of feeding behavior.

Notably, these appetite-suppressive effects of CB1R antagonists extend beyond food intake to drug-seeking behaviors. For instance, AM4113 (Balla et al., 2018) and AM6527 reduced alcohol intake across various paradigms, including drinking in the dark and intermittent access to alcohol (Raghav et al., 2024). Additionally, both AM4113 and AM6527 have been shown to suppress heroin-seeking behavior under FR and progressive (PR) schedules, as well as heroin and cocaine self-administration and reinstatement (AlKhelb et al., 2022; He et al., 2019; Soler-Cedeño et al., 2024). These findings suggest that CB1R antagonists consistently reduce operant responding for palatable food as well as for drugs of abuse, including alcohol, cocaine, and opioids.

One potential mechanism for these suppressive effects involves CB1R modulation of mesolimbic dopamine (DA) signaling. CB1Rs are predominantly located on presynaptic terminals throughout the mesolimbic system, including on glutamatergic afferents to the nucleus accumbens and on GABAergic interneurons in the basolateral amygdala (Covey & Yocky, 2021; Katona et al., 2001; Peters et al., 2021; Winters et al., 2012). These neuroanatomical substrates are likely critical for the motivational and consummatory behaviors suppressed by CB1R antagonism (Wenzel & Cheer, 2018) and these findings implicate a critical role for CB1R signaling in mesolimbic DA transmission. Supporting this, AM4113 was reported to reduce ethanol-induced DA release in the nucleus accumbens while decreasing alcohol intake without affecting locomotion or tastant preference (Balla et al., 2018). Similarly, CB1R antagonism has been shown to block THC-induced DA elevations in the accumbens (Tanda et al., 1997; Wenzel & Cheer, 2018). Therefore, the ability of CB1R antagonists to reduce motivated behavior likely reflects, at least in part, a reduction in mesolimbic dopamine transmission, highlighting the critical role of DA in mediating the behavioral effects of CB1 signaling.

Overall, AM6527 is a potent and orally bioavailable neutral CB1 receptor antagonist that exhibits high CB1 selectivity and improved pharmacokinetic properties relative to earlier compounds such as AM4113 and AM6545 (Sink et al., 2009). Prior studies have demonstrated that AM6527 reduces food-reinforced operant behavior in rats (e.g., Sink et al., 2009) and, more recently, suppresses ethanol consumption and seeking behavior in both male and female mice (e.g., Raghav et al., 2024). While its broad appetite-suppressing effects are well established, the specific behavioral processes disrupted by CB1 antagonism, or involvement of DA transmission, remain less understood. To address this, we employed a multi-level behavioral approach combining machine learning-assisted unsupervised behavioral classification by motion-sequencing (MoSeq), naturalistic reward consumption, FR and PR operant paradigms for palatable reward, and dopamine-targeting pharmacological manipulations. Together, this integrative strategy allows for a detailed dissection of how CB1R antagonism impacts spontaneous, consummatory, and motivational behaviors. Given its oral bioavailability, behavioral efficacy, and cleaner side-effect profile, AM6527 may represent a promising lead compound for the development of therapeutics targeting maladaptive motivation, including weight loss, obesity and other psychiatric and metabolic disorders.

## Methods

### Animals

Forty-eight adult male C57BL/6 mice were obtained from Jackson Laboratory and were housed in a colony maintained at 21 °C ± 2 and humidity (50 ± 20%) with a reversed 12-h light/dark cycle (lights off at 07:00 AM). Mice weighed between 22.1 and 26.7 g at the beginning of the study. Lab chow and water were available ad libitum. Mice that are trained in operant conditioning experiments were subjected to mild water deprivation by utilizing 1% citric acid in their daily water (Urai et al., 2021). All the behavioral sessions and drug testing were conducted during the dark phase of the light/dark cycle on weekdays. All experiments were approved by the Institutional Animal Care and Use Committee at McLean Hospital, which met the guidelines outlined in the NIH Guide for the Care and Use of Laboratory Animals.

### Pharmacological agents and selection of doses

**AM6527** was synthesized at the Northeastern University Center for Drug Discovery (Yohn et al., 2015)(Raghav et al., 2024) and was dissolved in 4% DMSO, 8% Tween-80, and 88% sterile saline (0.9%). DMSO/Tween/Saline mixture was used as a vehicle control. The lead time of AM6527 (30 minutes) and the dose range were chosen based on previous studies (Raghav et al., 2024). **SKF38393** (a D1 dopamine receptor agonist) was purchased from Tocris Bioscience (Cat. No. 0922) and dissolved in sterile saline (0.9%) solution, which was also used as vehicle. The lead time (20 minutes) and dose of SKF38393 were selected based on previous literature (Yohn et al., 2015) and pilot data. **Istradefylline** (an adenosine A2A receptor antagonist) was purchased from Cayman Chemical (Cat. No. 22958) and dissolved in 10% DMSO, 15% Tween-80, and 75% sterile saline (0.9%). The same solution was used for vehicle injections. Dose range and lead time (30 minutes) were based on published findings (Ecevitoglu et al., 2023) and pilot data. **GBR12909** (a dopamine transporter inhibitor) was obtained from Tocris Bioscience (Cat. No. 0421) and prepared in sterile saline (0.9%), which was also used as vehicle. The dose and lead time (30 minutes) were chosen based on prior studies (Yohn et al., 2016) and pilot data. All treatments were administered intraperitoneally (IP) at a volume of 10 ml/kg.

### Behavioral Procedures

#### Open Field Test

Spontaneous activity was assessed using a circular open field apparatus (radius: 42 cm). Mice were placed individually in the center of the arena and allowed to explore freely for 20 minutes. The arena was cleaned with 70% ethanol between subjects. The spontaneous behaviors were acquired through an Xbox One Kinect camera and analyzed by MoSeq (Wiltschko et al., 2015).

#### Naturalistic Reward Consumption Assay

To assess the effects on palatable fluid consumption, mice were individually housed and tested in their homecages using evaporated milk (Casa Solana, Houston, TX, USA). First, milk was introduced into the homecage to allow for initial exposure. The following day, mice received a single habituation injection of saline. Baseline fluid intake was then assessed on the subsequent day, during which mice received saline injections (IP) prior to testing. Drug testing began the following week and followed a within-subjects, counterbalanced design. On test days, mice received IP injections three hours into the dark cycle. Thirty minutes later, they were given access to pre-weighed bottles containing evaporated milk and water. After a 4-hour access period, bottles were removed, and fluid intake was measured.

#### Fixed Ratio 3 (FR3) Reinforcement Paradigm

Mice were trained on the FR3 nose poking paradigm for evaporated milk (Casa Solana, Houston, TX, USA) in 30-minute sessions (5 days/week, one session per day). Instrumental conditioning sessions were conducted in operant conditioning chambers for mice (15.2 x 17.8 x 18.3 cm; Med Associates, Fairfax, VT, USA), equipped with two nose-poke ports and a central dipper (0.2cc capacity). First, milk was introduced into the homecage to allow for initial exposure. Next, mice received one day of magazine training, during which evaporated milk was delivered on a variable intertrial interval (0–60 seconds, mean = 30 seconds) with each delivery accompanied by a 10-second dipper presentation. Mice were then trained on an FR1 schedule for up to one week, during which each correct nose poke triggered a 5-second dipper presentation. Next, mice were trained under an FR3 schedule for up to 2 weeks, with each reward delivery accompanied by a 3-second dipper presentation. During the following two weeks, mice continued FR3 training while receiving IP saline injections 2–3 times per week prior to drug testing. During testing, the number of head entries, correct nose pokes, and incorrect nose pokes were recorded.

#### Progressive Ratio (PR) Reinforcement Paradigm

Another cohort of mice was initially trained using the FR3 nose-poke paradigm for evaporated milk as described above. After a stable FR3 performance is observed within 1-2 weeks, they were transitioned to the PR schedule for another week in which the ratio was determined by the formula n =(5×e0.2i)-5 (Richardson & Roberts, 1996), and the program timed out after 3 minutes of no successful reward delivery to reinforce the breakpoint (maximum session length = 30 minutes). During the following two weeks, mice continued PR training while receiving IP saline injections 2–3 times per week prior to drug testing. During testing, the breakpoint (active nose poke time in minutes), the highest ratio achieved, the number of head entries, and correct nose pokes, and incorrect nose pokes were recorded.

### Experimental procedures

#### Experiment 1: Effect of AM6527 on spontaneous activity in the open field

The first cohort of mice (n = 16) received IP injections of vehicle or AM6527 (1.0, 3.0, or 10.0 mg/kg) twice per week in a within-subjects, counterbalanced design with at least 48 hours between treatments.

#### Experiment 2: Effect of AM6527 on naturalistic reward consumption

A second cohort of mice (n = 8) received vehicle or AM6527 (1.0, 3.0, or 10.0 mg/kg) twice per week in a within-subjects, counterbalanced design with at least 48 hours between treatments.

#### Experiment 3: Effect of AM6527 on FR3 operant conditioning for palatable reward

A third cohort of mice that were trained on the FR3 schedule (n = 8) was administered with vehicle or various doses of AM6527 (1.0, 3.0, or 10.0 mg/kg) once per week, following a within-subjects, counterbalanced design. When baseline sessions occurred the day after drug administration, behavioral performance generally returned to pre-treatment levels.

#### Experiment 4: The ability of dopaminergic drugs to reverse the effects of AM6527 on mice trained on FR3 schedule

The third cohort of trained mice (n = 6; one mouse was excluded due to health issues following completion of Experiment 3, and another due to a significant drop in baseline performance) was used for a drug reversal study. On drug treatment days, animals received two separate injections: one consisting of vehicle or AM6527 (the lowest dose from the previous experiment), and the other consisting of vehicle or one of three reversal compounds: SKF38393 (D1 receptor [D1R] agonist), istradefylline (adenosine A2A receptor [A2R] antagonist) or GBR12909 (dopamine transporter [DAT] blocker). The experiment employed a within-subject design, with each mouse receiving all five combined treatment conditions in a randomized order once per week: AM6527 vehicle + reversal agent vehicle (0.9% saline), 1.0 mg/kg AM6527 + vehicle, 1.0 mg/kg AM6527 + 5.0 mg/kg GBR12909, 1.0 mg/kg AM6527 + 0.5 mg/kg SKF38393, and 1.0 mg/kg AM6527 + 0.6 mg/kg istradefylline. When baseline sessions occurred the day after drug administration, behavioral performance generally returned to pre-treatment levels.

#### Experiment 5: Effect of AM6527 on PR operant conditioning for palatable reward

A fourth cohort of mice that were trained on PR schedule (n = 16) was administered with vehicle or various AM6527 (1.0, 3.0, or 10.0 mg/kg) once per week, following a within-subjects, counterbalanced design in which all animals received all treatments in a randomly varied order. When baseline sessions occurred the day after drug administration, behavioral performance generally returned to pre-treatment levels.

### Statistical Analysis

To assess spontaneous behaviors, trial videos were analyzed using MoSeq, a machine learning-assisted, unsupervised behavioral classification tool (Wiltschko et al., 2015), to extract and quantify sub-second behavioral syllables. Raw depth video files were first preprocessed to align and center the mouse in each frame. Principal component analysis (PCA) was then applied to reduce the dimensionality of the mouse’s pose dynamics, and a robust autoregressive hidden Markov model (ARHMM) was used to segment the behavioral syllables. These syllables represent recurring patterns of movement that are automatically identified and categorized by the model. Output metrics included syllable usage and duration, which were then used to compare behavioral patterns across drug doses, along with 2D speed of the animal per condition which is defined as the overall velocity during testing.

The rest of the behavioral data were analyzed using GraphPad Prism 10.0 (San Diego, CA, USA). First, data were assessed for normality and sphericity using the Shapiro–Wilk and Mauchly’s tests, respectively. Where violations of sphericity were detected, Greenhouse–Geisser corrections were applied. Mixed-effects models were used if needed to account for missing data. One-way repeated-measures analysis of variance (ANOVA) was used to assess the effects of AM6527 on syllable duration and usage during open field test, fluid intake during naturalistic reward consumption, FR3 performance, and PR performance during operant paradigms. Post hoc comparisons were conducted using Dunnett’s method for multiple comparisons, only comparing performance between the vehicle and various AM6527 doses. For reversal studies, a similar approach was used in which AM6527 + vehicle was compared with the other treatments. For studies utilizing PR operant performance, to evaluate differences between high and low performers on the task, mice were divided into performance groups based on a median split of vehicle-day nose poke counts. Using this grouping, factorial ANOVAs were conducted to analyze PR performance with the performance group included as a between-subjects factor. Significant interaction effects are reported, indicating that the impact of drug treatment varied depending on the baseline performance group. Statistical significance was defined as p < 0.05. All graphs display group means ± standard error of the mean (SEM).

## Results

### Experiment 1. MoSeq detects the alterations in spontaneous behaviors affected by AM6527

To assess the effects of AM6527 on spontaneous behavior, mice were tested in an open field and analyzed using MoSeq which detected changes in several different syllables of behavior upon administration of various doses of AM6527. The analysis revealed 47 unique syllables with both usage and duration. For the grooming face #1 syllable (Fig. 1A), a significant main effect of treatment was observed on usage (F(2.37, 31.54) = 5.52, p = 0.0063), with post hoc comparisons revealing increases at 3.0 and 10.0 mg/kg compared to vehicle (p = 0.0068 and 0.0073, respectively). Duration of this motif (Fig. 1D) was not significantly affected overall (F(2.39, 31.88) = 1.83, p = 0.17). Similarly, usage of grooming face #2 (Fig. 1B) was significantly increased by AM6527 (F(2.10, 28.00) = 7.51, p = 0.0022), with post hoc tests indicating higher usage at 10.0 mg/kg (p = 0.0032). The duration of grooming face #2 (Fig. 1E) was significantly increased (F(1.77, 23.61) = 7.80, p = 0.0033), with post hoc analyses showing elevations at both 3.0 and 10.0 mg/kg (p = 0.047 and p = 0.0091, respectively). AM6527 also increased expression of hind leg scratching (Figs. 1C and 1F), as indicated by both usage (F(1.95, 25.96) = 28.45, p < 0.0001) and duration (F(1.96, 26.06) = 16.78, p < 0.0001). Usage was significantly higher at 3.0 and 10.0 mg/kg (p = 0.0027 and p < 0.0001), and duration was elevated at those same doses (p = 0.0098 and p = 0.0001, respectively).

**Figure 1.**
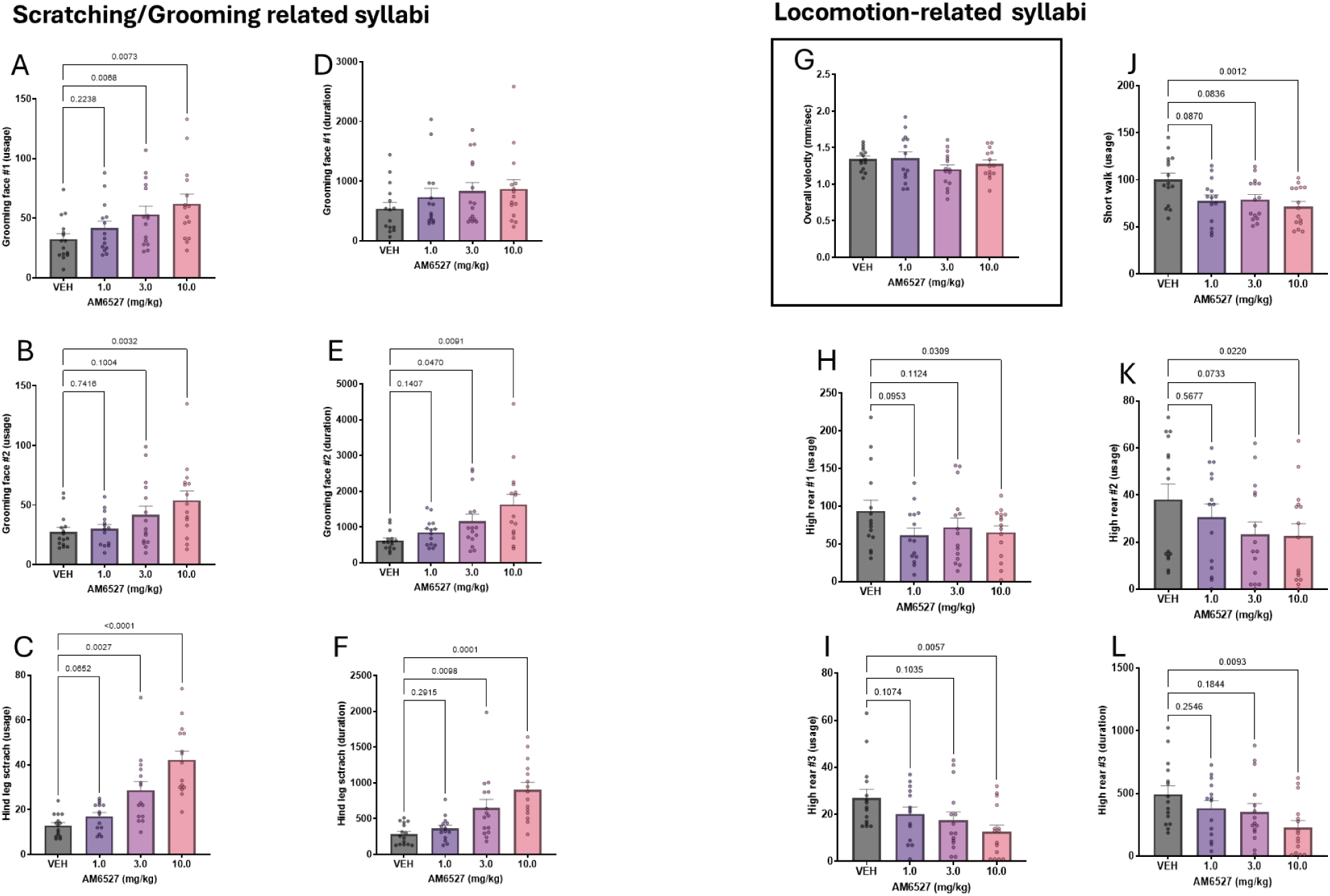
MoSeq analysis detects the alterations in spontaneous behaviors affected by AM6527. Usage frequency of grooming face #1 (A) was significantly increased at 3.0 and 10.0 mg/kg (**p < 0.05**), grooming face #2 (B) was significantly increased at 10.0 mg/kg (**p < 0.05**), and hind leg scratching (C) was significantly increased at both 3.0 and 10.0 mg/kg (**p < 0.05**). Duration of grooming face #1 was not significantly affected (D; **p > 0.05**), while grooming face #2 (E) was significantly increased at 3.0 and 10.0 mg/kg (**p < 0.05**), and hind leg scratching duration (F) was significantly increased at both 3.0 and 10.0 mg/kg (**p < 0.05**). (G) AM6527 had no significant effect on overall locomotor velocity (**p > 0.05**). Usage of high rear motif #1 (H), high rear motif #2 (I), and high rear motif #3 (J) were significantly reduced at 10.0 mg/kg (**p < 0.05**), as was the duration of high rear motif #3 (L; **p < 0.05**). Usage of the short walk syllable (K) was also significantly reduced at 10.0 mg/kg (**p < 0.05**), with non-significant trends at 1.0 and 3.0 mg/kg (**p > 0.05**). Data are presented as mean ± SEM.

In terms of locomotion, AM6527 did not influence the overall velocity (mm/sec; Fig.1G; F(1.774, 23.65) = 1.620, p = 0.2200) or affected general motor function (unpublished data). Yet, MoSeq detected several changes in locomotion related syllables. The short walk syllable (Fig. 1J) was significantly reduced in usage (F(2.49, 33.14) = 6.08, p = 0.0034), with post hoc tests revealing a significant decrease at 10.0 mg/kg (p = 0.0012), and trends toward reduction at 1.0 and 3.0 mg/kg (p = 0.087 and 0.084). Three distinct high rear motifs were also affected. For the first high rear syllable (Fig. 1H), usage was significantly reduced (F(2.37, 31.64) = 4.22, p = 0.0187), with a post hoc reduction at 10.0 mg/kg (p = 0.0309). The second high rear variant (Fig. 1K) showed a similar effect (F(2.74, 36.52) = 3.49, p = 0.0285), with a decrease at 10.0 mg/kg (p = 0.0220). For the third high rear motif (Figs. 1I and L), both usage (F(2.65, 35.30) = 6.33, p = 0.0022) and duration (F(2.83, 39.55) = 4.92, p = 0.0061) were decreased at 10.0 mg/kg, with reduced usage (p = 0.0057) and duration (p = 0.0094).

### Experiment 2. AM6527 reduces naturalistic reward consumption of milk but does not influence water intake

Milk intake in the homecage was significantly affected by AM6527 treatment (F(3, 21) = 10.89, p = 0.0002, η² = 0.609, Fig. 2A). Post hoc comparisons revealed a significant reduction at 10.0 mg/kg (p < 0.0001), while lower doses (1.0 and 3.0 mg/kg) did not differ significantly from vehicle (p = 0.5855 and p = 0.0855, respectively). In contrast, the water intake in the homecage (Fig. 2B) was unaffected by AM6527 (F(3, 27) = 1.18, p = 0.334).

**Figure 2.**
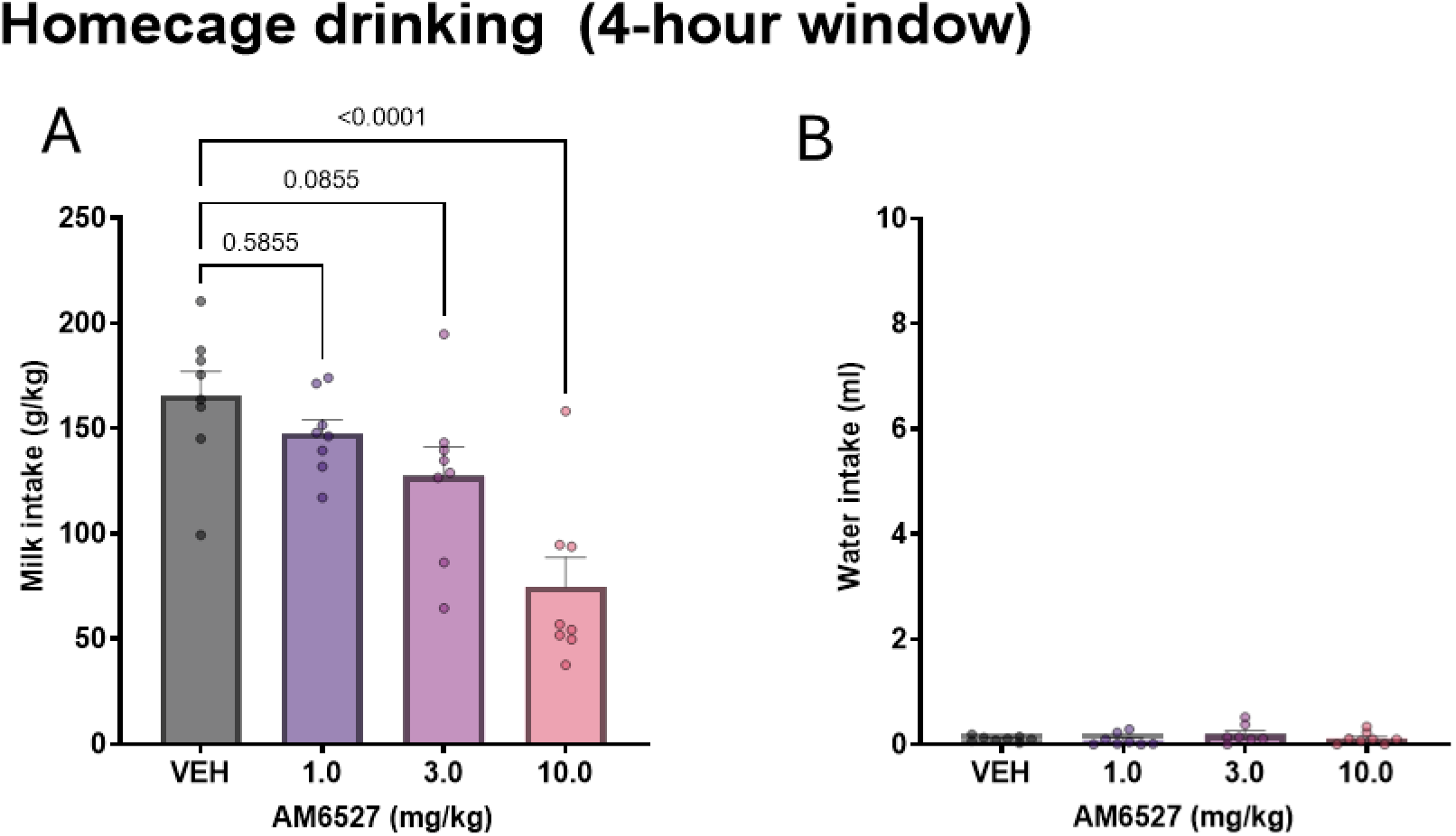
AM6527 reduces naturalistic reward consumption of milk but does not alter water intake. (A) Milk intake (g/kg) was significantly decreased at 10.0 mg/kg AM6527 (**p < 0.05**), with no significant effect at lower doses (**p > 0.05**). (B) Water consumption (ml) was not significantly affected by AM6527 at the tested dose range (**p > 0.05**). Data are presented as mean ± SEM.

### Experiment 3. AM6527 reduces operant responding for palatable reward under FR3 operant schedule

To assess the effects of AM6527 on operant behavior, mice were tested on an FR3 schedule for evaporated milk. Center port head entry counts (Fig. 3A) were significantly affected by AM6527 treatment (F(3, 21) = 6.91, p = 0.0021, η² = 0.497). Dunnett’s post hoc tests revealed reduced the number of head entries at 1.0 mg/kg (p = 0.0232), 3.0 mg/kg (p = 0.0180), and 10.0 mg/kg (p = 0.0007) compared to vehicle. AM6527 also produced a robust, dose-dependent suppression of nose poke responding (Fig. 3B; F(3, 21) = 30.98, p < 0.0001, η² = 0.816). Significant reductions were observed at all doses tested (1.0, 3.0, and 10.0 mg/kg; all p < 0.0001).

**Figure 3.**
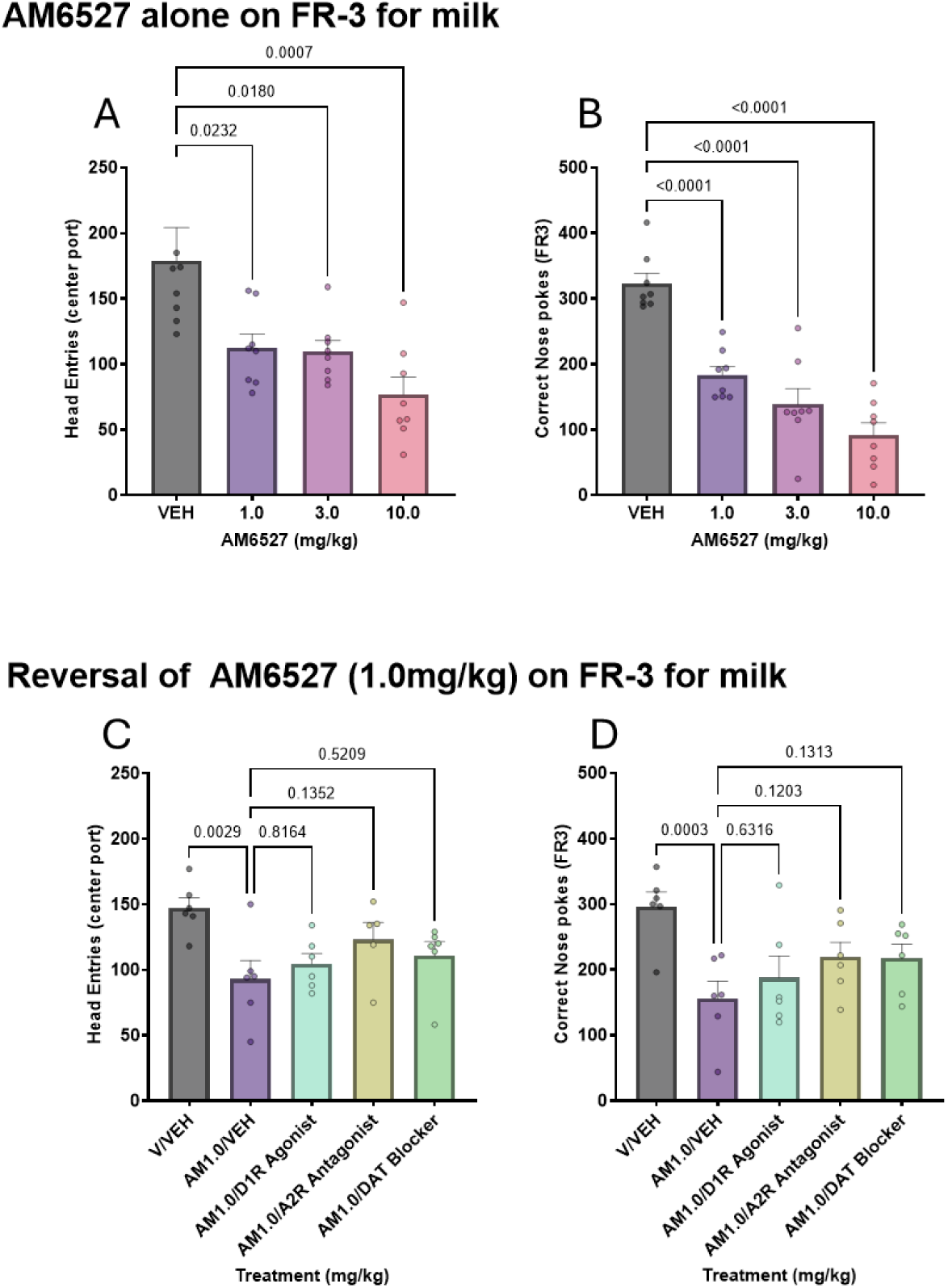
AM6527 suppresses operant responding for palatable reward under FR3 schedule and is not fully reversed by dopaminergic manipulations. AM6527 significantly reduced the number of center port head entries (A) and nose poke at all doses tested (**p < 0.05**). Co-administration of D1R agonist, A2AR antagonist, or DAT blocker did not significantly reverse AM6527-induced reductions in the number of head entry (C) or nose pokes (D; all **p > 0.05**), although A2AR and DAT treatments produced intermediate but non-significant effects. Data are presented as mean ± SEM.

### Experiment 4. Effects of AM6527 on FR3 performance is not fully reversed by dopamine-based interventions

AM6527 at 1.0 mg/kg significantly reduced FR3 performance compared to vehicle, and this effect was evaluated in combination with selective dopaminergic and adenosinergic agents. The number of center port head entries showed a significant overall treatment effect (F(4, 19) = 4.61, p = 0.0091; Fig. 3C). Post hoc comparisons revealed that AM6527 alone (AM1.0/VEH) produced a significant reduction compared to vehicle-treated animals (V/VEH; p = 0.0029), similar to Experiment 3. Co-administration of a D1R agonist (p = 0.816), A2AR antagonist (p = 0.135), or DAT blocker (p = 0.521) failed to significantly restore head entry counts, although A2AR and DAT manipulations showed numerically intermediate values. Nose poke responding was also significantly altered (F(4, 20) = 6.68, p = 0.0014; Fig. 3D). Compared to V/VEH, AM1.0/VEH produced a robust suppression of responding (p = 0.0003), similar to Experiment 3. None of the reversal conditions significantly differed from AM1.0/VEH: D1R agonist (p = 0.632), A2AR antagonist (p = 0.120), or DAT blocker (p = 0.131), indicating partial but statistically nonsignificant mitigation of the suppressant effect.

### Experiment 5. AM6527 reduces PR schedule performance for palatable reward with effects modulated by baseline performance

To assess the motivational effects of AM6527, mice were tested on a PR schedule of reinforcement. Analyses were conducted first across all subjects and subsequently based on response performance under vehicle condition.

Repeated measures ANOVA revealed a significant main effect of treatment on center port head entry counts (F(3, 45) = 3.36, *p* = 0.0268, η² = 0.183; Fig. 4A). Post hoc comparisons indicated a significant reduction at the 10.0 mg/kg dose relative to vehicle (*p* = 0.0101), whereas the 1.0 and 3.0 mg/kg doses did not differ significantly (*p* = 0.6166 and *p* = 0.6997, respectively). Nose poke responding was also significantly modulated by AM6527 (F(2.02, 30.35) = 14.70, *p* < 0.0001, η² = 0.495; Fig. 4B). Compared to vehicle, AM6527 significantly reduced nose poke counts at both 3.0 and 10.0 mg/kg (*p* = 0.0096 and *p* = 0.0002, respectively). A trend-level reduction was observed at 1.0 mg/kg (*p* = 0.0779). Analysis of the highest ratio achieved demonstrated a robust effect of treatment (F(3, 48) = 14.53, *p* < 0.0001, η² = 0.476; Fig. 4C). All doses of AM6527 significantly reduced the highest ratio relative to vehicle (*p* = 0.0320, 0.0012, and < 0.0001 for 1.0, 3.0, and 10.0 mg/kg, respectively). In contrast, breakpoint duration did not reach statistical significance (F(3, 48) = 2.33, *p* = 0.0865, η² = 0.127; Fig. 4D).

**Figure 4.**
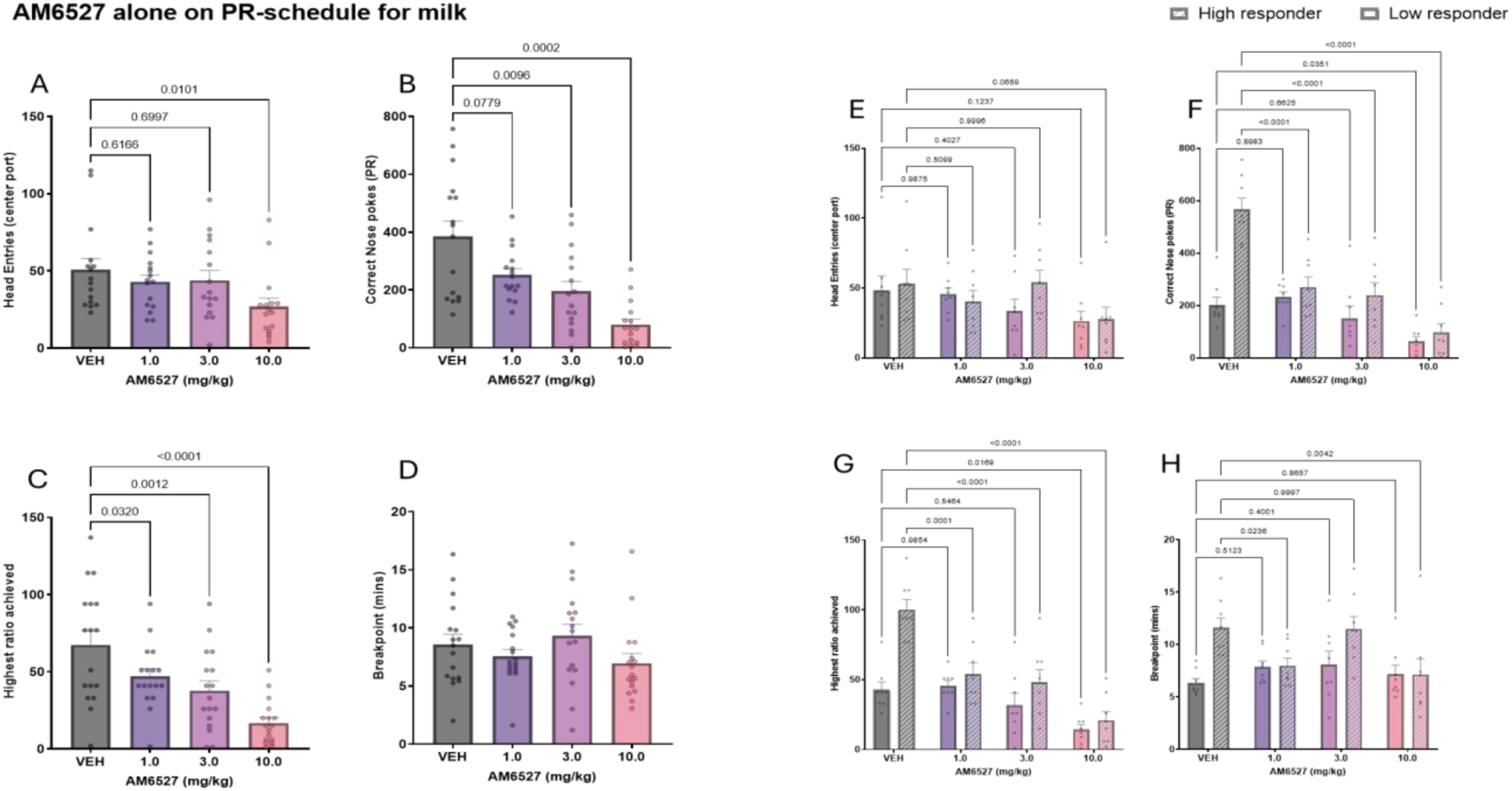
AM6527 impairs motivation on PR schedule performance, with effects modulated by baseline performance. In the full cohort, AM6527 significantly reduced center port head entry counts (A) at 10.0 mg/kg (**p < 0.05**), nose poke counts (B) at 3.0 and 10.0 mg/kg (**p < 0.05**), and highest ratio achieved (B) at all doses tested (1.0, 3.0, and 10.0 mg/kg; **p < 0.05**). (D) Breakpoint duration was not significantly affected by AM6527 at any dose (**p > 0.05**). (E) Stratified analysis by baseline performance (median split) showed a significant main effect of treatment on the number of head entries (**p < 0.05**), but no significant main effect of response group or treatment × group interaction (**p > 0.05**); within-group post hoc comparisons were not significant for any dose (**p > 0.05**). (F) Nose poke counts were significantly reduced in high responders at 1.0, 3.0, and 10.0 mg/kg (**p < 0.05**), while low responders showed a significant reduction only at 10.0 mg/kg (**p < 0.05**). (G) Highest ratio achieved was significantly reduced in high responders at all doses (1.0, 3.0, and 10.0 mg/kg; **p < 0.05**) and in low responders at 10.0 mg/kg only (**p < 0.05**). (H) Breakpoint duration was significantly reduced in high responders at 1.0 and 10.0 mg/kg (**p < 0.05**), with no significant changes in low responders (**p > 0.05**). Data are presented as mean ± SEM.

To better understand individual variability in drug sensitivity, we performed a post hoc stratification based on vehicle PR performance. Mice were divided into high and low responder groups using a median split of total nose pokes under vehicle conditions. This classification was then used as a between-subjects factor in two-way repeated measures ANOVAs across each dependent measure. For the number of head entries to the center port (Fig. 4E), the analysis revealed a significant main effect of treatment (F(3, 42) = 3.36, *p* = 0.0276), but no main effect of response group (*p* = 0.4653) or treatment × group interaction (*p* = 0.4111). When analyzed separately, neither group showed statistically significant differences across doses, though the 10.0 mg/kg dose showed a trend toward reduction in both groups (*p* = 0.124 in low responders, *p* = 0.066 in high responders). For correct nose poke counts (Fig. 4F), there were significant main effects of treatment (F(3, 42) = 22.11, *p* < 0.0001), group (F(1, 14) = 34.95, *p* < 0.0001), and a robust treatment × group interaction (F(3, 42) = 8.57, *p* = 0.0001). Post hoc tests indicated that all doses significantly reduced nose poke counts in high responders (all *p* < 0.0001), while in low responders, only the 10.0 mg/kg dose had a significant effect (*p* = 0.0351). The highest ratio achieved (Fig. 4G) also showed significant main effects of treatment (F(3, 42) = 20.32, *p* < 0.0001), group (F(1, 14) = 29.74, *p* < 0.0001), and a treatment × group interaction (F(3, 42) = 5.71, *p* = 0.0023). High responders displayed dose-dependent reductions at all doses (*p* <0.0001), whereas low responders only showed a significant reduction at 10.0 mg/kg (*p* = 0.0169). Finally, breakpoint duration was also significantly affected by treatment (F(3, 42) = 3.15, *p* = 0.0348), group (F(1, 14) = 7.03, *p* = 0.0190), and their interaction (F(3, 42) = 3.89, *p* = 0.0154). Among high responders, both 1.0 and 10.0 mg/kg doses significantly reduced breakpoint time (*p* = 0.0236 and *p* = 0.0042, respectively), while no significant effects were observed in low responders (*all p* > 0.05).

## Discussion

The present findings demonstrate that AM6527, a neutral CB1 receptor antagonist, produced dose-dependent changes in spontaneous behavior, operant responding, and palatable reward consumption in male mice. Although the antagonist did not affect overall velocity and general behaviors, it altered sub-second behavioral syllables associated with grooming and locomotion, as identified through machine learning-assisted MoSeq analysis. In the naturalistic reward consumption assay, milk intake was significantly reduced at 10.0 mg/kg, while water intake remained unaffected and no changes were observed in the preference. Under an FR3 operant schedule, AM6527 suppressed the number of head entries and nose pokes at all doses, and these effects were not significantly reversed by dopaminergic agents, although A2A antagonist and DAT blocker did show a trend in reversing the effects of AM6527. In the PR task, AM6527 decreased motivation for reward, as indicated by reductions in nose poke counts and highest ratio achieved, with more pronounced effects in high-responding animals.

Using MoSeq, we identified dose-dependent alterations in spontaneous behavior, characterized by a reduction in locomotion-related syllables (e.g., walking, rearing) and an increase in grooming- and scratching-related syllables. These effects emerged most clearly at the highest two doses, and notably occurred without any changes in overall velocity, highlighting the sensitivity of MoSeq in detecting discrete, behaviorally specific changes. Given that these shifts occurred within brief behavioral motifs (400 ms syllables across 20 minutes), MoSeq enables quantification of subtle motor and ethological changes that might be missed by conventional scoring approaches. For instance, the increased grooming and scratching behaviors observed here are consistent with prior observations using AM4113 and other related CB1R antagonist, which have also been associated with excessive grooming and scratching (Hodge et al., 2008; Jarbe et al., 2006, 2008; Sink et al., 2010). Together, these findings suggest that AM6527 alters spontaneous behavior in a manner consistent with other CB1R antagonists, and that MoSeq provides a powerful tool for detecting these ethologically relevant changes induced by neuroactive and psychoactive drugs.

The naturalistic reward consumption assay further demonstrated that AM6527 reduces consummatory behavior. A significant decrease in palatable reward intake was observed only at the highest dose, confirming that CB1R antagonism suppresses intake even in the absence of a work requirement. Importantly, milk preference over water remained unchanged at all doses, indicating that hedonic valuation of the palatable fluid was preserved and that the observed effects are unlikely to reflect taste aversion. This interpretation is supported by prior studies showing that AM6527 does not induce gaping responses in feeding paradigms (Sink et al., 2009). Similarly, our previous work found that mice continued to consume highly palatable super saccharin solutions and exhibited no reductions in water intake, suggesting preserved preference (Raghav et al., 2024) across both studies.

In the operant paradigms, AM6527 reduced effort for palatable reward across both FR3 and PR schedules. In the FR3 task, AM6527 suppressed operant responding for palatable reward, and in the PR schedule, reduction was particularly evident in high responders, based on a median split of baseline vehicle performance. These results are consistent with prior findings that CB1 antagonists reduce operant responding across both low- and high-effort schedules, including FR1 and FR5 (Salamone et al., 2007), suggesting that the suppressive effects of CB1 blockade on reward seeking generalize across response demands. Additionally, the absence of change in overall locomotor velocity in the open field, as demonstrated by MoSeq analysis, indicates that these behavioral effects were not attributable to gross motor deficits. Instead, MoSeq identified selective, small, but significant reductions in locomotion-related syllables and increases in grooming-related syllables at the sub-second scale, suggesting an ethologically specific shift in behavior rather than sedation.

Since CB1 receptors modulate synaptic transmission at key nodes in the mesolimbic system as discussed above, influencing DA release and regulating glutaminergic and GABAergic inputs to the nucleus accumbens and basolateral amygdala, we investigated whether pharmacologically increasing dopaminergic signaling could reverse AM6527’s behavioral effects. As illustrated in Fig. 3, the FR3 performance suppression by AM6527 for palatable reward was not fully reversed by dopaminergic compounds. These dopaminergic rescue experiments revealed partial and inconsistent reversal of AM6527’s effects by D1 receptor activation (SKF38393) and D2 receptor modulation by A2A receptor antagonism (istradefylline), or DAT blocker (GBR12909). These findings suggest that AM6527’s behavioral effects are mediated by DA-dependent and -independent pathways. This indicates that CB1R antagonism may influence motivation through both dopaminergic and alternative mechanisms, such as non-dopaminergic modulation of limbic or cortical inputs.

In addition to suppressing motivation for palatable stimuli, AM6527 has also been found to mitigate consumption and self-administration of various drugs of abuse. Findings from our recent study (Raghav et al., 2024) demonstrate that AM6527, at the same doses used here (1, 3, and 10 mg/kg), robustly reduced ethanol intake in male and female C57BL/6J mice across multiple paradigms, including drinking-in-the-dark, intermittent access, and operant chain schedules. These effects were dose-dependent, long-lasting, and did not exhibit tolerance across repeated dosing, similar to other CB1R antagonists such as AM4113 (Balla et al., 2018), consistent with our findings of persistent suppression across both effort-based (FR3, PR) and naturalistic reward consumption assays. Interestingly, AM6527 appeared more effective in suppressing motivation for ethanol and palatable rewards than in prior drug self-administration studies, where significantly higher doses (10, 20, and 30 mg/kg) were required to inhibit cocaine and opioid intake (Soler-Cedeño et al., 2024). Yet, these findings build around previous ideas (Salamone et al., 2007) that CB1R antagonists could be therapeutically valuable for disorders involving heightened motivation for food and drugs of abuse.

Collectively, the findings from this study align with prior studies using AM4113, AM6527, and AM6545, which have shown reductions in operant and consummatory behaviors without impairing locomotion or inducing conditioned taste aversion (Chambers et al., 2007; Randall et al., 2010; Sink et al., 2008, 2009, 2010). Our data significantly advances the current understanding of AM6527’s actions by using MoSeq analysis to precisely capture behavioral changes and by investigating underlying mechanisms through dopaminergic rescue attempts. This suggests that compounds like AM6527 are promising therapeutic options for disorders involving heightened reward sensitivity such as obesity and alcohol use disorder, as they offer the advantages of avoiding negative side effects or significant motor impairment often associated with dopaminergic interventions.

